# Porcupine: Rapid and robust tagging of physical objects using nanopore-orthogonal DNA strands

**DOI:** 10.1101/2020.03.06.981514

**Authors:** Kathryn Doroschak, Karen Zhang, Melissa Queen, Aishwarya Mandyam, Karin Strauss, Luis Ceze, Jeff Nivala

## Abstract

Molecular tagging is an approach to labeling physical objects using DNA or other molecules that can be used in cases where methods like RFID tags and QR codes are not suitable. No molecular tagging method exists that is inexpensive, fast and reliable to decode, and usable outside a lab setting to create or read tags. To address this, we present Porcupine, an end-user molecular tagging system that features DNA-based tags readable within seconds using a portable nanopore device. Porcupine’s digital bits are represented by the presence or absence of distinct, nanopore-orthogonal DNA strands, which we call molecular bits (molbits). We classify molbits directly from the raw nanopore signal, avoiding basecalling. To extend the tag’s shelf life, decrease readout time, and make tags robust to environmental contamination, molbits are prepared for readout during tag assembly and can be stabilized by dehydration. The result is an extensible, real time, high accuracy tagging system that includes a novel approach to developing nanopore-orthogonal barcodes.

**One sentence summary:** Porcupine lets end-users label physical objects with custom DNA tags, without requiring a lab to create or read tags, and offers rapid readout using nanopore sequencing.

## Introduction

DNA-based information storage continues to gain momentum with the emergence of high-throughput DNA sequencing (sequencing-by-synthesis) and synthesis (array-based synthesis); these technologies enable vast amounts of text and visual (*1–3*) information to be encoded, stored, and decoded. DNA data storage offers unique advantages over mainstream storage methods like magnetic tape and hard disk drives, including higher physical density and longer retention lifetimes (*4*). This is particularly useful for archival storage, where current access speeds and read/write costs are not an issue. The development of portable, real-time sequencing (nanopore) (*5*), together with new methods that simplify the modular assembly of pre-defined DNA sequences (*6*), creates additional opportunities for rapid writing and on-demand readout. This compelled us to revisit an early application of DNA-based information storage: molecular tagging (*7*).

*Molecular tagging* uses physical molecules to encode an identifier for a physical object, analogous to a radio-frequency identification (RFID) tag or Quick Response (QR) code in the digital world. An ideal molecular tagging system should be inexpensive and reliable, with fast readout and user-controlled encoding and decoding from end-to-end. Molecular tags can be easily used in situations where RFID tags and QR codes are not suitable; for example, labeling and tracking commodities that are too small, flexible, or numerous to attach a sticker or electronic tag, especially to detect counterfeits. Other promising applications include tracking and establishing provenance for higher value items, or covertly exchanging private information like encryption keys and digital wallets. Although several molecular tagging solutions exist, none is fully usable – from creation to readout – outside of a laboratory setting, or without involving a third party. These constraints significantly increase tag read and write latency, decrease tag confidentiality, and ultimately limit custom application development.

To address these issues, we designed and built Porcupine, a molecular tagging system that uses synthetic DNA-based tags. Although DNA is typically considered expensive for reading and writing, Porcupine lowers the cost by providing a fixed library of 96 pre-synthesized, nanopore-orthogonal DNA fragments, which we call molecular bits (molbits). Molbits represent digital bits through presence or absence instead of 1s and 0s (Fig. 1a); allowing new, arbitrary tags that end-users can create for custom applications by simply mixing molbits to produce a molecular tag. Molecular tags are then read out quickly using a portable, low-cost sequencing device (Oxford Nanopore Technologies MinION; Fig. 1b). Typically, raw nanopore signal must first be converted back to a DNA sequence in a computationally expensive process called base-calling, but we classify molecular tags directly from raw nanopore signal, forgoing basecalling. Raw signal classification is often used for DNA and RNA sample demultiplexing, which also uses DNA barcodes (*8, 9*); Porcupine drastically increases the number of barcodes by custom designing them to produce unique ionic current signatures. Error correction is also added to the tag to resolve decoding errors, similar to electronic message transmissions systems. Molbits are prepared for readout (sequencing) prior to tag application and can be stabilized by dehydration, a new approach that extends tag shelf life, decreases decoding time, and reduces contamination from environmental DNA. The result is a highly accurate real time tagging system that includes a novel approach to developing nanopore-orthogonal barcodes. These barcodes, and the methods we use to develop them, are extensible; they can be used both within Porcupine to tag physical objects and beyond this system for other molecule-level tagging needs like sample multiplexing for nanopore sequencing

**Figure 1:**
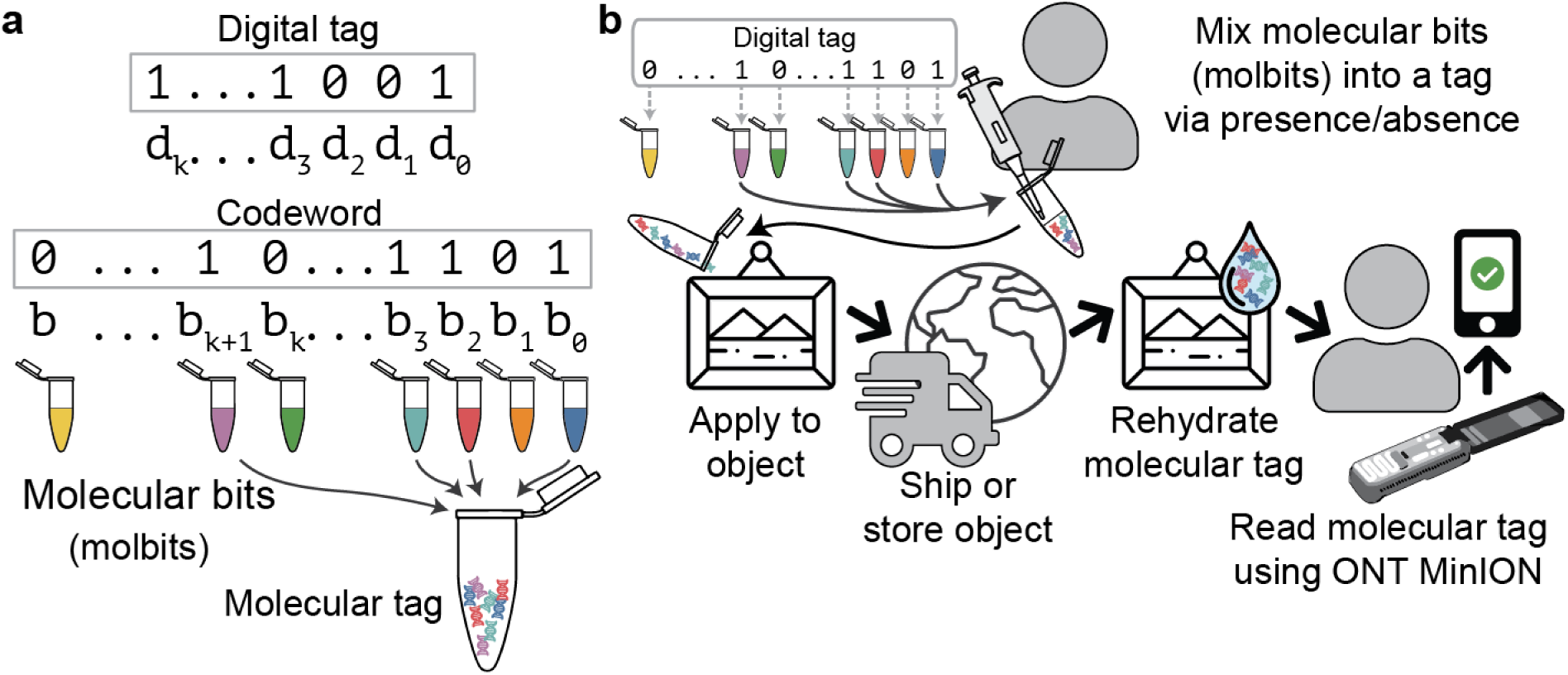
Creating a molecular tag using the Porcupine molecular tagging system. (a) Porcupine’s encoding scheme. A digital tag is converted into a codeword to add additional bits for error correction. Each codeword bit is assigned to a unique molbit, where 1s and 0s are represented by the presence or absence of individual molbits in the molecular tag mixture. (b) A user first defines a digital tag as a binary 96-bit number, and pipettes 1-bits into the molecular tag. The tag is applied to an object, which is then shipped or stored. To read the tag, it is rehydrated and loaded directly onto Oxford Nanopore Technologies’ MinION device. Software then identifies the tag, either with or without knowledge of the original tag.

## Results

To develop Porcupine, we first defined an individual molbit as a DNA strand that combines a unique barcode sequence (40 nt) with a longer DNA fragment selected from a set of sequence lengths that we pre-determined (Fig. 2a). To make assembly of molbits simple and modular, we designed them to be compatible with Golden Gate Assembly by incorporating a short single-stranded overhang. To increase classification accuracy and decrease computation time, we further optimized them to avoid basecalling. Thus, for the barcode region, the objective was to produce a large set of sequences that could generate unique ionic current signatures (“squiggles”) to promote unambiguous classification.

**Figure 2:**
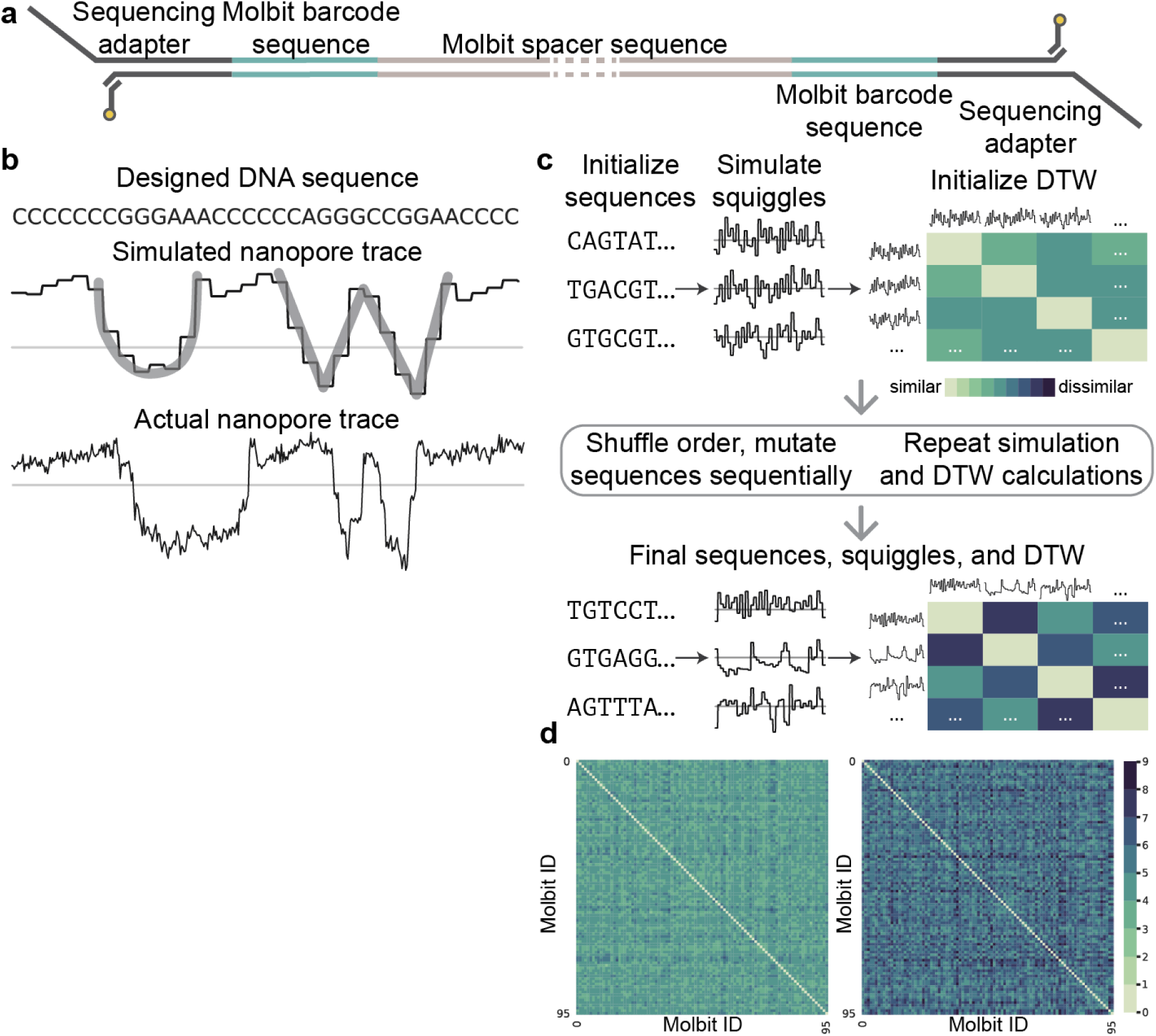
Molbit design scheme. (a) Molecular bit (molbit) structure. The molbit sequence is attached to a spacer sequence via Golden Gate assembly to achieve a minimum length for sequencing and provide an additional encoding channel. Since the sequencing adapter is attached to both ends, the strand can be sensed from either direction. (b) The letters “UW” depicted visually in nanopore raw data (as opposed to encoded in the sequence contents). From top to bottom, the shown sequence was simulated using Scrappie and sequenced on the ONT Min-ION, demonstrating the viability of using simulations for designing intentional, arbitrary raw signal shapes. (c) Evolutionary model workflow. Each round of evolution begins with a set of sequences, their simulated squiggles, and pairwise Dynamic Time Warping (DTW) distances. The sequence order is randomized, and sequences are mutated one at a time, verifying DTW improvement (minimum and mean) after each attempt. (d) Dynamic time warping (DTW) scores before (left) and after (right) 31 iterations of the evolutionary model. After initialization, the minimum DTW similarity was 2.9 (mean 4.2 +/- 0.4), and after evolution the minimum was 4.2 (mean 5.8 +/- 0.8).

To model the predicted ionic current signature for arbitrary DNA sequences, we used Scrappie squiggler, a tool that converts sequences of bases to ionic current via a convolutional model. For example, to demonstrate Scrappie’s ability to accurately model real nanopore squiggles, we designed a DNA sequence that appears as the letters UW in squiggle space (Fig. 2b), with high visual similarity to the simulated squiggle (except for noise). Scrappie’s output also let us compute the signal similarity of two sequences quantitatively using dynamic time warping (DTW) as the distance measure. We used this approach inside an evolutionary model designed to make barcodes as separable as possible (Fig. 2c).

To produce a set of 96 orthogonal molbit barcode sequences, we initialized the evolutionary model using 96 random or pre-seeded starting sequences (see Methods). We perturbed each sequence independently in random order by mutating two adjacent nucleotides simultaneously at a random location. If the mutated sequence failed to improve the minimum and average DTW similarities between itself and all other sequences, we reversed the mutation and attempted again for the same sequence. We also restricted sequence similarity and free energy of the sequences to avoid labeling ambiguities and secondary structure (see Methods). Using this method, we began with a set of starting sequence that had a minimum DTW similarity of 2.9 and mean 4.2 +/- 0.4 and achieved a final minimum of 4.2 and mean 5.8 +/- 0.8 after 31 rounds of evolution (Fig. 2d), representing a 44.8% and 38.1% improvement in the minimum and mean, respectively.

Given the set of 96 designed molbit barcodes, we wanted to increase the number of available molbits without requiring additional barcode design or synthesis. To do this, we inserted a DNA fragment between the barcode regions as a spacer sequence, which can be set to different lengths as an additional encoding channel. Thus, since each molbit consists of the unique combination of a molbit barcode plus a specific spacer sequence length, adding another length effectively adds an additional 96 molbits. Length works as an additional encoding channel because even without basecalling, the length of nanopore signals can be easily distinguished; the signal length is roughly proportional to the DNA fragment length. We tested this concept using two spacer lengths (400 and 1600 nucleotides) and found that simple signal length binning was sufficient for decoding. The median signal length for the 400 and 1600 nt strands was 5768 and 16968, respectively, and a cutoff at 9800 gave 91.0% accuracy, where most errors were caused by long strands misidentified as short strands (Fig. 3a). For remaining experiments, we focused on a single spacer length.

**Figure 3:**
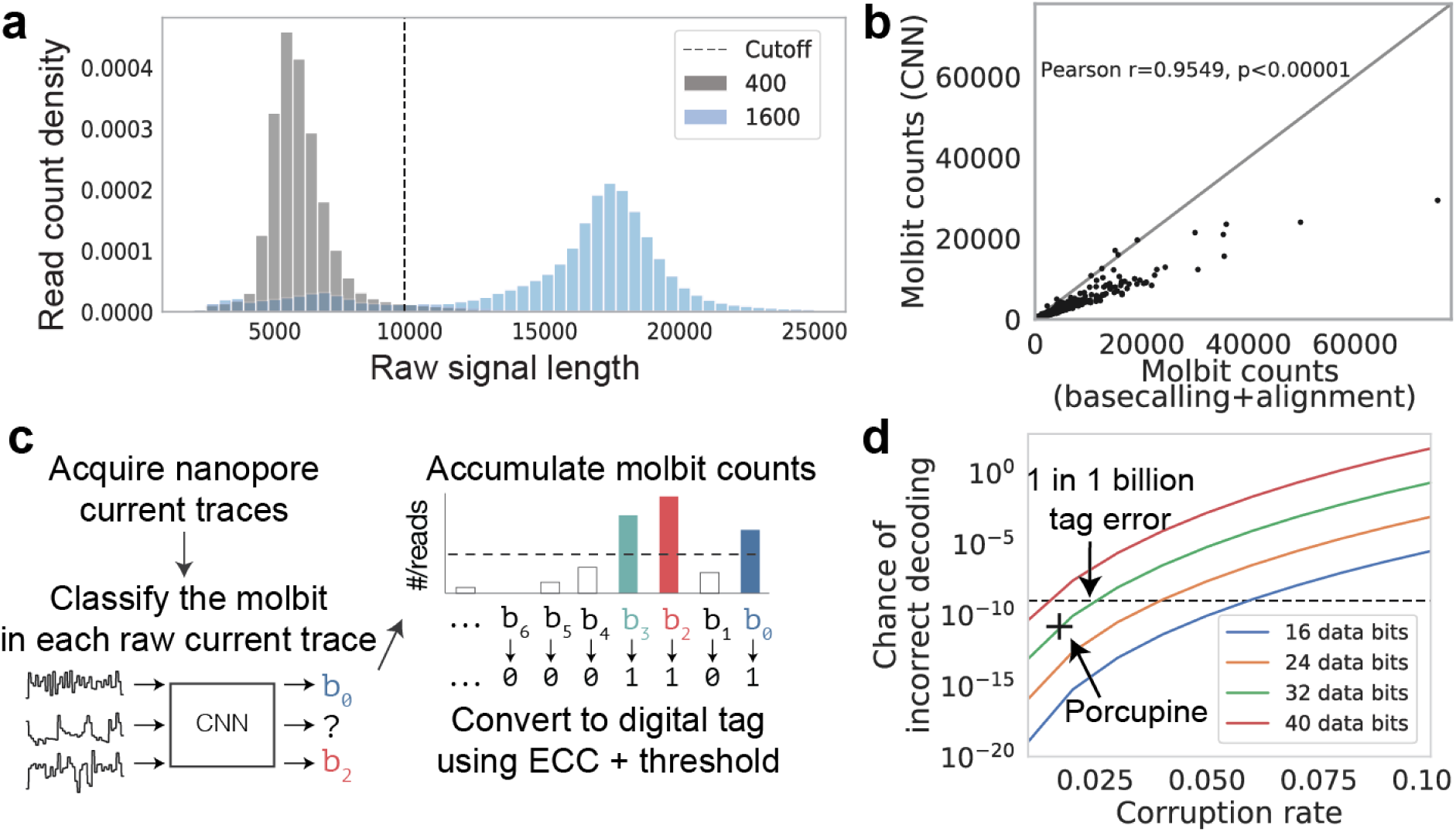
Molbit classification and tag decoding results. (a) Distribution of raw nanopore signal lengths for two different length DNA sequences (400 and 1600 nt), using kernel density estimation. (b) Correlation of read counts for each molbit, demonstrating consistency in molbit occurrences between training and testing runs (Pearson correlation, *p* = 3.1 × 10^−26^, *r* = 0.84). Counts were first normalized within each run and normalized again after combining runs for either training or testing. (c) Tag decoding workflow, with error correcting codes (ECC). After acquiring nanopore current traces from a standard sequencing run, the molbit in each trace is identified using the CNN (confidence ≥ 0.9). Successfully identified molbits are accumulated and converted into binary using a threshold for presence. This threshold is varied as error correction is carried out multiple times, accepting the binary digital tag with the most likely correct decoding. (d) Chance of incorrect tag decoding as a function of the bit corruption rate and number of data bits. This chance increases exponentially as the corruption rate and number of data bits increase linearly. The dashed line represents the goal of “1 in 1 billion tags incorrect,” and the “+” marks Porcupine’s chance of incorrect decoding.

After designing the molbits and acquiring a theoretical understanding of their separability, we developed a convolutional neural network (CNN) model to accurately classify them directly from real raw nanopore data (see Supplementary Figure 1). Note that molbits do not need to be segmented or otherwise isolated from the raw data, rather, the CNN uses only the first portion of the each molbit since the molbit barcode is located at the beginning of the strand. We gathered training data by dividing the 96 molbit barcodes into 6 sequencing sets, with each molbit appearing once; optimizing the sets for maximum sequence separability to improve labeling (see Methods); and running each set on the MinION. We assigned labels to each molbit read using traditional basecalling methods and a modified semilocal Smith-Waterman sequence alignment (*10*), using only high-confidence alignments (see Methods). For test data, we divided the 96 molbits into 2 sequencing sets, with each molbit appearing once, and gathered the data as described for training. The model was trained for 108 iterations, with a final training, validation, and test accuracy of 99.93%, 97.70% and 96.96%, respectively, compared to sequence-derived labels. However, in real world decoding, all reads are classified, not just those that pass basecalling and sequence alignment. We found that the CNN was consistently able to confidently classify a larger portion of the reads (97% of reads in the test set) than basecalling plus alignment (75.1% of reads in the test set), revealing a large portion of reads that we could not easily validate. We reasoned that if the occurrence of each molbit was proportional between basecalling and the CNN, that the CNN was likely not making spurious calls, but was perhaps performing better on the raw signal data. This seems to be the case as read counts correlate extremely well between the two methods (see Supplementary Figure 2). As a result, “accuracy” only reflects the model accuracy and does not necessarily measure each molbit’s error rate or the overall chance of decoding the tag incorrectly.

We next composed actual molecular tags. We assigned each molbit a unique position in a binary tag, allowing each 1 or 0 to represent the presence or absence of a specific molbit over the course of a single tag sequencing (decoding) run. To determine presence or absence, we used CNN-classified read counts for each molbit. Ideally, 0 bits would have zero reads, and 1 bits would have nonzero reads. However: two factors complicated our setting this threshold to determine bit presence: (1) nonzero read counts for molbits not present in an experiment, and (2) significant variations in counts for molbits present in an experiment, which we found to be up to 20-30x in our training and test sets (see Supplementary Text). Fortunately, these variations were consistent when we compared the ratios of molbit counts in the training and test data (Fig. 3b). We accounted for this variation by scaling all read counts by a fixed vector based on these ratios. Thresholding and scaling read counts reduced our per-bit error rate from 2.96 +/- 1.83% to 1.70 +/- 1.68%, a 42.6% reduction.

Since a reliable tagging system should have a very low chance of incorrect decoding (e.g., 1 in 1 billion), we decided to further reduce our overall tag decoding error rate by including error correcting codes (ECCs) as part of our tag design (Fig. 3c). The simplest non-ECC method for encoding information in these tags is a naive 1:1 mapping between digital bits and molbits; however, with this method, even a single bit error makes the tag unrecoverable (i.e., produces an incorrect decoding). In our system, bits are set to 1 or 0 using a threshold for presence or absence on the read counts, meaning that any 0-bits above this threshold are instead flipped to 1, and vice versa. ECCs reduce the possibility of unrecoverable tags despite the relatively high per-bit error rate by reserving a smaller number of bits for the digital message and creating a codeword by projecting this message into a larger space with greater separability. This allows more bits to be flipped before the message is decoded incorrectly. To encode the digital message, we simply multiply the message by a binary matrix of random numbers, known as a random generator matrix (See Supplementary Text). The number of bits reserved for the ECC depends on the application’s error tolerance and the per-bit error rate (Fig. 3d). As the error rate increases, the chance of incorrect decoding increases exponentially. Thus, the number of bits for the message must be chosen carefully. We chose a message size of 32 bits, which at an error rate of 1.70% produces a 1.6 × 10^−11^ chance of incorrect decoding and permits ∼4.2 billion total unique tags.

Next, as a proof-of-principle for our tagging system, we demonstrated end-to-end tag encoding and decoding of the acronym MISL, short for Molecular Information Systems Lab (Fig. 4a). We began by encoding MISL into binary using ASCII, which uses 8 bits for each character, for a total of 32 bits, and we multiplied this bit vector by the generator matrix to produce a 96-bit codeword. The molecular tag was then prepared as explained previously, with one modification for lab efficiency (see Methods). Once the molecular tag was assembled, it was prepared for sequencing and read out using an ONT MinION. We then identified the molbits from the raw data using our trained CNN classifier, accumulated a count for each molbit, and rescaled these counts (as explained above) to accommodate systematic read count variances. We then decoded the tag by binarizing the counts with varying thresholds and finding the nearest valid codeword. The earliest correct decoding occurred less than 7 seconds after sample loading (109 molbit strands observed), demonstrating reliable encoding and decoding of a 32 bit message in only a few seconds using a portable sequencing instrument.

**Figure 4:**
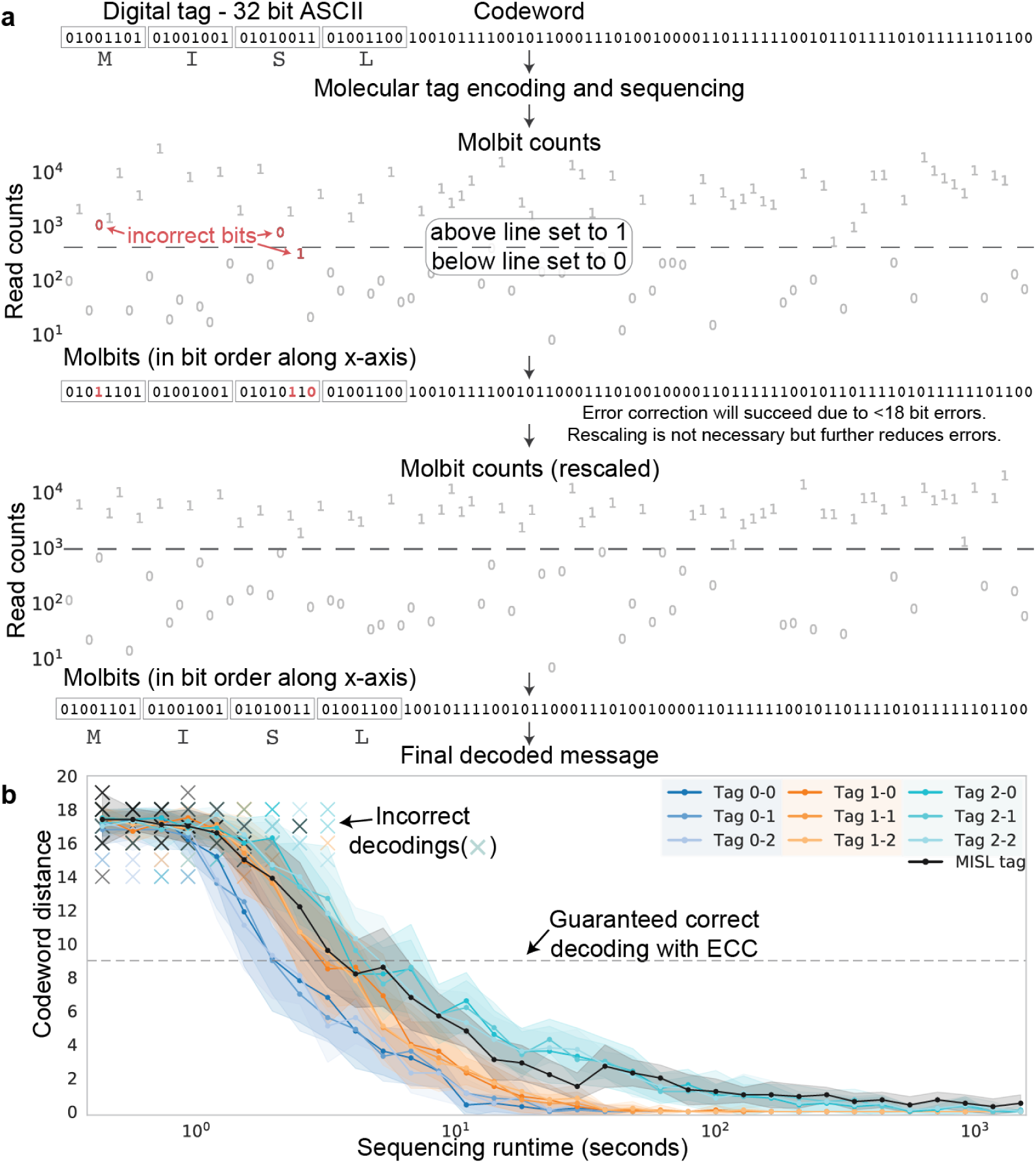
End-to-end data flow for the acronym “MISL”. (a) Encoding began by converting “MISL” to ASCII (32 bits). The digital tag was then multiplied by the 32×96-bit generator matrix, producing a 96-bit codeword. The codeword was converted to a molecular tag, stored, and sequenced for 35 minutes. When sequenced, molbits were identified and accumulated into a single count for each molbit. Bit errors are calculated with respect to the closest codeword, not necessarily the correct codeword. The dashed line represents the optimal threshold for binarizing these counts, which produced three incorrect bits. Rescaling the counts according to known read count variation reduced these errors, in this case eliminating errors entirely. (b) Minimum tag decoding distance as a function of sequencing runtime. Decoding occurred without a priori knowledge of the correct tag. Bit errors are capped by the distance between tags, which is at least 18 in the chosen ECC. Incorrect decodings are marked with an X. At each time point, reads were sampled 10x at an average of 10k reads per minute.

Finally, to robustly estimate correct decoding time under these conditions, we simulated tags using our original two test runs and one additional random tag. To generate a sufficient amount of simulation data, we composed synthetic datasets representing new codewords that were generated with the same number of 1-bits as the original tag datasets. The original observed molbits were then randomly assigned to the new bit ordering: 1-molbits in the original tag were assigned new 1-molbit labels in the synthetic tag. To do this, we sampled reads without replacement, accumulated a count for each molbit, and decoded as above, with 10 repetitions per simulated run time. This simulation impacts the error rate both positively and negatively due to the distribution of distances between codewords in the ECC. We found that some tags could be successfully decoded with only a few seconds of sequencing data, and all tags after just 10-15 seconds (Fig. 4b).

In summary, Porcupine offers a new method for molecular tagging based on the presence or absence of synthetic DNA sequences that generate explicitly unique nanopore raw current traces. By directly manipulating segments of nanopore raw current and keeping unique sequences short, we reduce synthesis costs for the end user and produce visually unique nanopore current traces, enabling high accuracy decoding. The speedy decoding time means our system can decode using newer technologies such as the Flongle, a cheaper, single-use flowcell produced by ONT with a quarter of the pores of the MinION in as little as 1-3 minutes. In addition, tags can be prepared for sequencing at the time of tag creation and then shelf-stabilized by dehydration. This further reduces readout time while maintaining tag quality (see Supplementary Figure 3). In the future, more bits can be acquired by adding more insert lengths, by extending the length of the unique region to allow more variation between molbits, or by combining barcode regions serially. Furthermore, a generative model for molbit design may be a natural next step, especially if a dramatically larger number of molbits is desired, because the evolutionary model computations scale exponentially with the number of desired bits.

One limitation of Porcupine is the variation in molbit counts. Although decoding remains robust to bit errors, more stability in read counts would increase the amount of information we could encode by reducing the number of bits required for error correction. Without resolving this variation, the number of bits can still be substantially expanded by adding more insert lengths to take advantage of the modular system design.

From a computational perspective, we note that basecalling is getting faster and more reliable; however, raw signal classification is fundamentally a simpler problem than basecalling. Such classifiers can be trained with comparatively minimal data and expanded to non-traditional sensing; for example, our molbit approach could be easily extended to include non-standard bases for additional security, but standard basecalling would not be possible without extensive amounts of training data.

## Acknowledgments

We thank additional members of the Molecular Information Systems Lab for feedback on this work and also express our gratitude to Sergey Yekhanin (Microsoft) for helpful discussions about error correction codes.

## Funding

This project was funded in part by DARPA under the Molecular Informatics Program and gifts from Microsoft.

## Author Contributions

K.D. participated in conceiving the idea and design/analysis of experiments, data curation, formal analysis, methodology, software, validation, visualization, writing - original draft, and writing - review & editing. K.Z. participated in investigation, validation, and writing - review & editing. A.M. contributed software for error analysis, as well as writing - review & editing. M.Q. contributed the ECC software and visualization, as well as writing review and editing. K.S. participated in investigating the ECC strategy, in the design/analysis of experiments and supervised the work. L.C. participated in conceiving the idea and design/analysis of experiments and supervised the work. J.N. participated in conceiving the idea and design/analysis of experiments, supervised the work, and contributed to writing original draft and writing review & editing.

## Competing Interests

A provisional patent has been filed by the University of Washington covering aspects of this work. JN is a consultant to Oxford Nanopore Technologies. KS is a Microsoft employee.

## Data and Materials Availability

Code is available at https://github.com/uwmisl/Porcupine. Data is available at https://github.com/uwmisl/Porcupine-data.

## Materials and Methods

### Molbit barcode design algorithm

To produce nanopore-orthogonal squiggles, we used an evolutionary model. First, we initialized the model using a set of starting sequences. Fully random sequence initialization works fairly well; however, we chose to initialize using a previous iteration of molbit barcodes, which produced slightly better results than random initialization after the molbits were further perturbed. Starting sequences and their corresponding simulated nanopore squiggles are shown in Supplementary Figure 4.

During mutation, we placed constraints on the sequences to ensure that they can be easily synthesized, assembled, and measured using the ONT MinION. If a mutated sequence does not fulfill these constraints, the mutation is reversed and attempted again. There are two types of constraints: (1) those that affect only a single sequence (independent constraints), and (2) those that impact the relationship between one sequence and all others (dependent constraints). Independent constraints require each sequence to be within a range of allowed GC content (30-70% GC), have a maximum folding potential (−8 kcal/mol) as calculated using NUPACK’s MFE utility (*11*), exclude the BsaI cut site sequence (GGTCTC), and have a maximum homopolymer length of five for A/T and four for C/G. Dependent constraints require a minimum sequence dissimilarity, calculated using a local variant of the Smith-Waterman (SW) algorithm (*10*) (≤ 15 SW score; cost function +1 match, - 1 mismatch, - 8 gap); and a minimum squiggle dissimilarity, calculated by simulating the sequence’s nanopore squiggle using the Scrappie squiggler and computing the dynamic time warping similarity (*12*) for all squiggles vs. the new squiggle.

At the start of each round of sequence evolution, sequence order is randomized. Each sequence is mutated sequentially in this random order. The mutation is introduced by simultaneously modifying two adjacent nucleotides, in a random location. If the new sequence fails to fulfill the preceding constraints, we undo the mutation and try again until a maximum number of tries (100, arbitrarily), at which point we proceed to the next sequence. Next, we recalculate sequence similarities with respect to the new candidate sequence. If any sequences are too close to the new sequence, we undo the mutation and try again. Next, the nanopore squiggle is simulated for the new sequence using Scrappie squiggler We recalculate the dynamic time warping similarity for all squiggles vs the new squiggle, and, if any squiggles are now too close, we undo the mutation and try again. If the new mutation improves both the minimum and the average DTW dissimilarity between all squiggles, it is accepted; if not, the mutation is reversed and reattempted. Evolution ends when the optimization begins bouncing between just two sequences. At this point, the process has produced a local minimum as the result of a series of random incremental improvements, so further improvements may be gained only by significantly perturbing these final sequences. We show final sequences and their corresponding simulated nanopore squiggles in Supplementary Figure 5.

### Molbit classification model

We identify individual molbits using a classification model, which takes raw nanopore signals as input and outputs the molbit ID with an associated confidence. The model consists of a 5-layer CNN, followed by two fully connected layers with 50% dropout, and a final fully connected layer with softmax as the output layer. Each of the 5 CNN layers is identically structured, including a 1D convolutional layer with Relu activation, average pooling, and then batch normalization. We show a diagram of the model with exact parameters for each layer (e.g., kernel size) in Supplementary Figure 1a.

Ideally, we we would build a training dataset by sequencing each of the 96 molbits separately. However, due to cost, we instead divided the 96 molbits into 6 runs of 16 molbits each. We constructed these sets to have a high predicted distance between the molbits within a set, meaning the most similar and easily confused molbits were not sequenced together for training data acquisition.

We assigned training labels using basecalling (Guppy version 3.2.2 with GPU acceleration) followed by Smith-Waterman sequence alignment (cost function: +1 match, - 1 mismatch, - 8 gap) against the full set of 96 molbits. We considered any SW score ≥ 15 to be a well-aligned match. As a quality measure, we also examined how many of these reads were labeled with one of the 16 possible molbits. An average of 98.7% +/- 2.1% of well-aligned reads belonged to the true set of molbits across all training runs, indicating high quality labels.

After labeling the training data, we balanced the dataset by allowing a maximum of 6000 reads occurrences for each molbit, with a total of 274,667 reads used for training. To pre-process the raw signal, we rescaled the signal using a Median Absolute Deviation method modified from Oxford Nanopore Technologies’ Mako classification tool, trimmed the signal to remove the variable-length stalled signal characteristic to the beginning of sequencing reads, and finally truncated the signal to the first 3000 data points.

We split the training data 85%/15% to produce training and validation sets and trained the model for 109 iterations, with a final maximum training accuracy of 99.94% and validation accuracy of 97.78%. Confusion matrices for testing and validating the final model are shown in Supplementary Figure 6a.

We acquired and labeled testing data in the same manner as the training data, using 2 new sequencing runs, each containing a unique half of the molbits. Performance on these test sets was 98.1% and 95.7% for labeled data. (See Supplementary Figure 6b for confusion matrices.) We washed and reused the flowcell from test set 1 for test set 2, which potentially contributed to a small portion of the errors present due to DNA carryover between the runs. We show read counts for these two test runs in Supplementary Figure 7, noting when a molbit was possibly present from the previous run.

### Experimental methods

We purchased forward and reverse strands of the 40 nt unique barcode sequences from Integrated DNA Technologies (IDT). Reverse strands contained a 5’ GATG overhang and a 3’ dA-tail. We annealed forward and reverse strands by mixing them equimolar in 0.5M PBS, boiling them at 94C for two minutes and then allowing them to cool at room temperature. To generate the insert spacer, we amplified an arbitrary 400 nt portion of plasmid pCDB180 (https://www.addgene.org/80677/) by PCR, using primers we designed to add BsaI cut sites to the ends of the amplified product.

To assemble the molbits, 600 ng of the desired annealed barcodes (at equimolar concentrations) and 600 ng of the spacer were ligated together using NEB’s Golden Gate Assembly Kit. We prepared molbits for sequencing using ONT’s Ligation Sequencing Kit (SQK-LSK109) following the kit protocol (we skipped the “DNA repair and end-prep” step because molbits were already dA-tailed) and ONT’s Flow Cell Priming Kit (EXP-FLP001). Molbits were sequenced on a R9.4.1 MinION flow cell with bulk FAST5 raw data collection enabled on MinKNOW.

We dehydrated tags after nanopore adapter ligation by mixing the molbits with 1% trehalose dihydrate solution and lyophilizing the sample. To sequence, we rehydrated the lyophilized sample in nuclease-free water and carried out sequencing prep described above.

## Supplementary Text

### Evaluating read count variation

As described in the main text, the largest source of bit errors in Porcupine can be attributed to the unusually large but seemingly consistent read count variance. Since each molecular tag is composed of equal proportions of each present molbit, the resulting read counts for each molbit after sequencing should be approximately equal; but instead, we measured significant variation in read counts. We first checked for obvious sequence-related issues by correlating quantifiable sequence characteristics, including GC content, minimum free energy (MFE), and homopolymer length, against read counts normalized as described in the main text. Evidence shows GC bias in genomic bacterial nanopore sequencing data on the MinION (*13*), based on non-stochastic bias introduced by the basecaller, not the sequencing device. With respect to MFE, we reasoned that although we controlled for MFE in the design phase, correlation between folding energy and read counts could still explain some variation. Additionally, we considered that the inclusion of long homopolymers could also have been associated with a reduction in read counts since imprecise calls of homopolymer length can potentially cause poor sequence alignment and therefore artificially low read counts (*14*). As shown in Supplementary Figure 8a, correlations between these various measures and normalized read counts were quite low (Pearson correlation for GC: p = 0.31, r=-0.10; MFE: p=0.33, r=0.09; homopolymer length: p=0.82, r=0.02). Thus, these rough measures could not explain the source of read count variance.

Moving beyond these initial sequence bias analyses, we reasoned that read count variation errors can be introduced at four different points in our system: (1) DNA synthesis, (2) strand assembly and tag combination, (3) sequencing, or (4) analysis/labeling.

(1) To check whether there was a problem with our original DNA order, we re-ordered 15 molbits representing the five lowest, average, and highest read count molbits. When sequenced, the read counts of the re-ordered sequences were similar in proportion to their previous counts (Supplementary Figure 8b). Thus, if the source of variability is caused by a synthesis error, which we believe is unlikely, it is at least reproducible or possibly dependent on an unexplored aspect of sequence content.

(2) To evaluate the strand assembly and tag combination step, we examined potential secondary structure in individual molbits, specifically the same molbits from the re-synthesized tests. During the Golden Gate strand assembly step, double stranded DNA is separated into single strands, potentially enabling secondary structure formation. Sequences were screened for minimum free energy during the evolutionary modeling phase, but the screening did not cover all forms of secondary structure, particularly stem-loop structures with short stems and large loops. Some secondary structure can be seen across all three categories (see Supplementary Figure 9a). It is possible that secondary structure could have lowered read counts, but we do not believe that it could have caused excessive read counts.

Additionally, we reasoned about potential experimental variation due to human error. When combining tags, some variation can be explained by pipetting. Here, we reasoned that up to 2-3x variation could be explained by this factor (bubbles, calibration, improperly depressed pipette, etc.). Also, if a bit were omitted, it would cause a near-zero or very low read count, but if a bit were accidentally added, it alone would not cause such a dramatic overabundance of read counts. Thus, human error of this type could cause a low read count but likely not an overly high one.

(3) It is possible the systematic sequencing errors contribute to molbit read differences, however, the strong correlation between basecalling and CNN-based molbit classification makes this less likely. Future experiments using a different sequencing platform (e.g. Illumina or Sanger), could help to resolve this question.

(4) We do not believe basecalling or labeling is a significant contributor to the problem. With respect to basecalling, an average of 93.3% of reads could be basecalled with high confidence (Q-score ≥ 9) throughout all runs, leaving just ∼7% room for variation (compared to 200-300% variation overall).

Also note that the molbit counts for labels assigned via basecalling and alignment have high correlation with those assigned by the CNN (Pearson *p* < 10^−5^, r=0.9998, Supplementary Figure 8c). Although there is certainly bias in that figure since basecalled labels are used to train the CNN, this high correlation does reduce the possibility that either method is introducing a significant number of spurious calls.

Ultimately, our method was designed to be robust to these types of errors; however, we still sought to reduce them since any reduction in bit errors can exponentially reduce the overall decoding error rate.

### Error correction

To improve the system’s resilience we overlay an error correcting code. The error correcting code maps the original messages into a higher dimensional space that provides a greater distance between any two messages. In this higher dimensional space a message must accumulate many errors before it is decoded incorrectly, allowing us to reduce the chance of incorrect decoding for a given fixed error rate.

The error correcting code protocol has two stages: encoding and decoding. We use a random linear code, which consists of a fixed, randomly chosen *n* × 96 generator matrix that encodes an *n*-bit message via a simple and efficient matrix multiplication. The resulting 96-molbit code-word may accumulate errors during creation, storage, or retrieval. However, we can decode the 96-molbit codeword back to the *n*-bit message with high probability using brute-force nearest neighbor decoding.

## Supplementary Figures

**Supplementary Figure 1:**
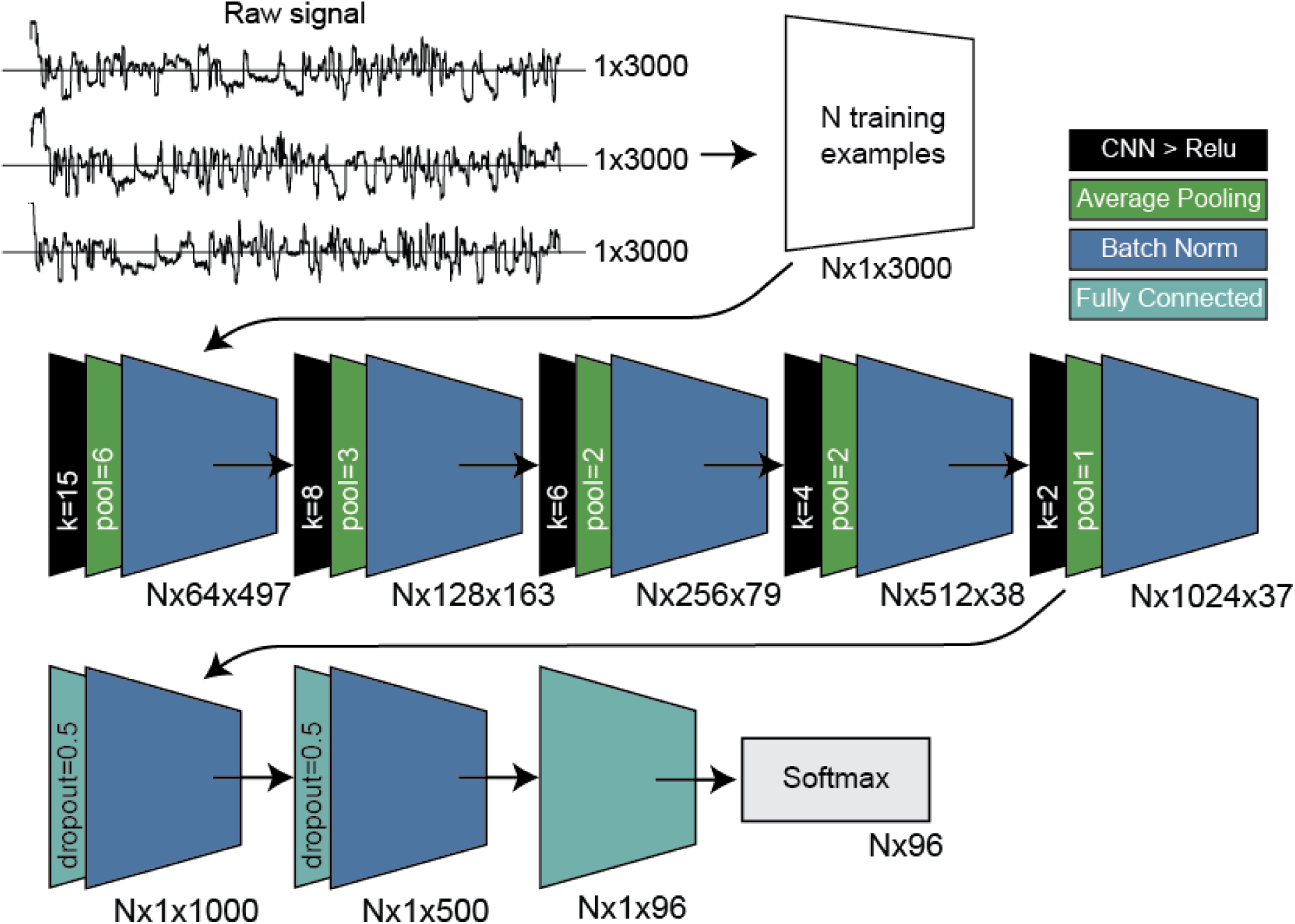
Molbit classification model and data flow. Input data includes raw nanopore signal, which was rescaled using a Median Absolute Deviation method modified from Mako, trimmed to remove stalled signal characteristic to the beginning of sequencing reads, and truncated to the first 3000 data points in the signal time series. Rescaled training data then passes through a 5-layer CNN followed by 2 fully connected layers with dropout and a final fully connected layer with softmax as the output layer. In the CNN layers, k is the kernel size and pool is the average pooling kernel size.

**Supplementary Figure 2:**
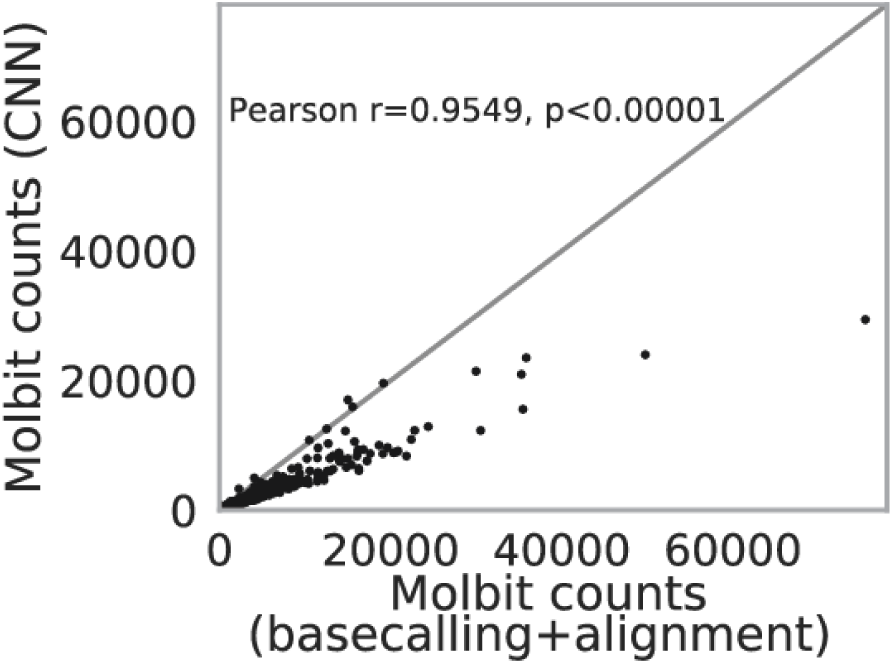
Comparison of molbit counts from sequence data (labeled using base-calling plus alignment) vs. signal data (labeled using CNN).

**Supplementary Figure 3:**
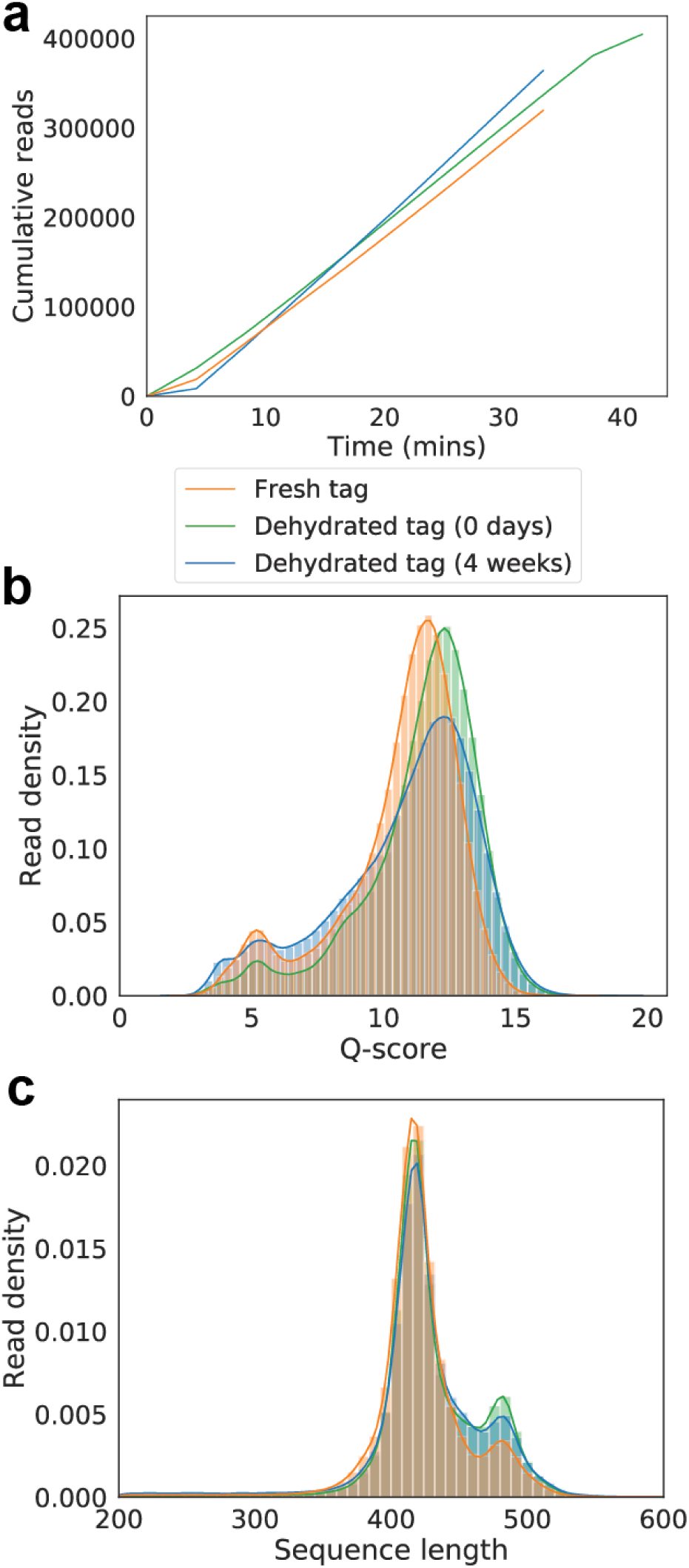
Quality of fresh versus dehydrated tags. (a) Cumulative sequencing read output over time. (b) Distribution of PHRED quality scores for all reads after basecalling using Guppy 3.2.2 (GPU version). (c) Distribution of sequence lengths after basecalling.

**Supplementary Figure 4:**
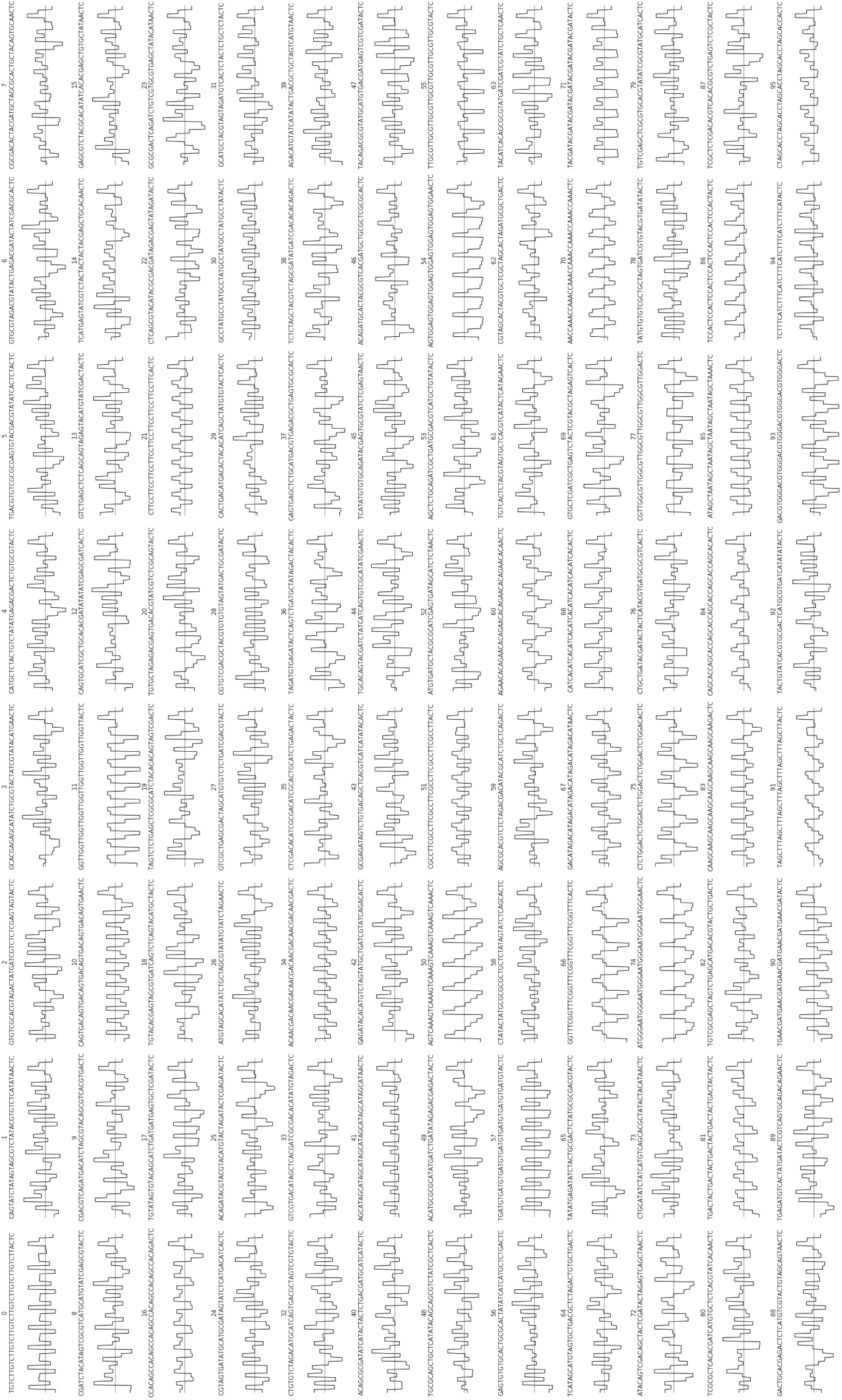
All 96 molbit sequences initialized before beginning the design phase using the evolutionary model.

**Supplementary Figure 5:**
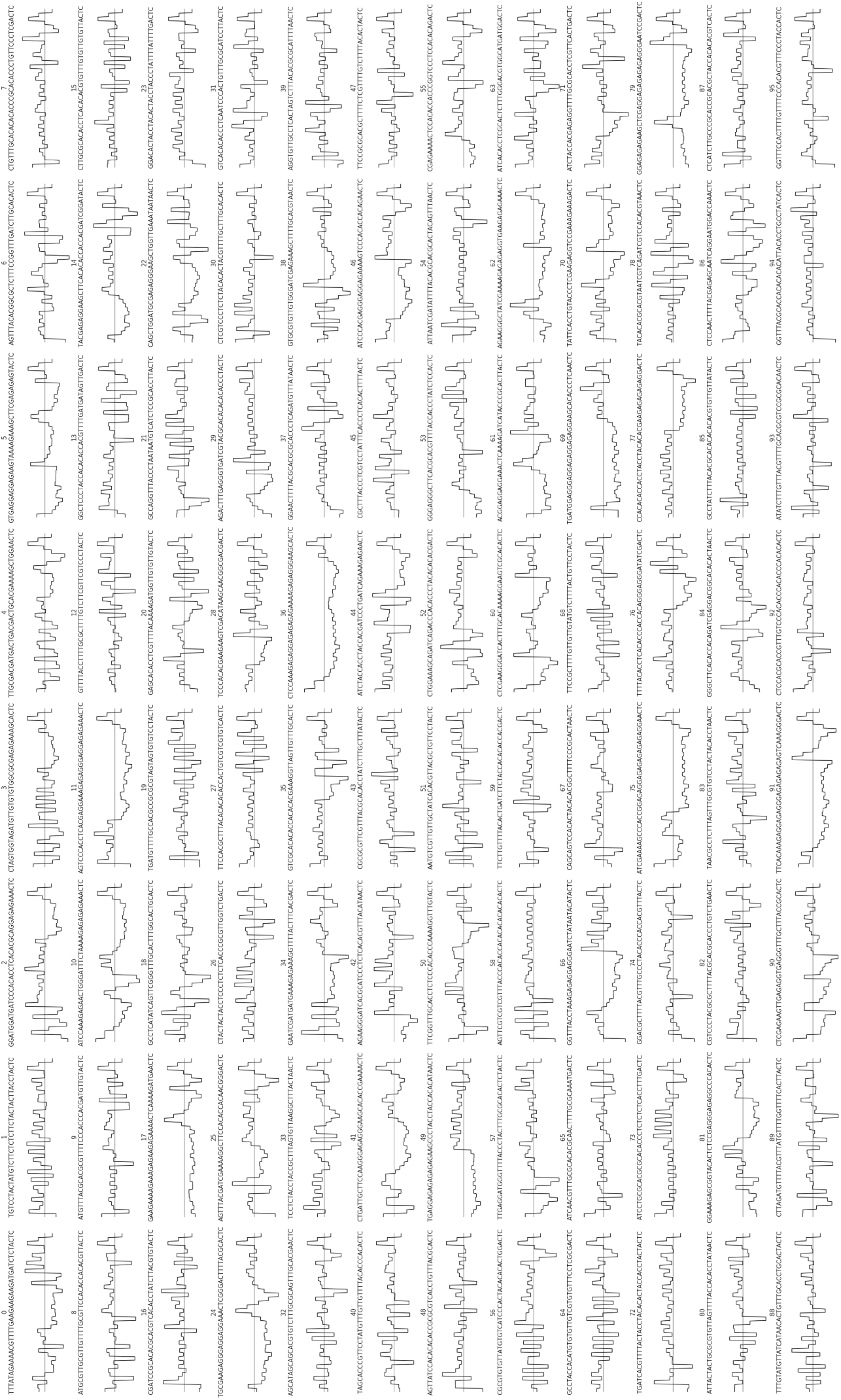
All 96 molbit sequences after 31 iterations of the evolutionary model.

**Supplementary Figure 6:**
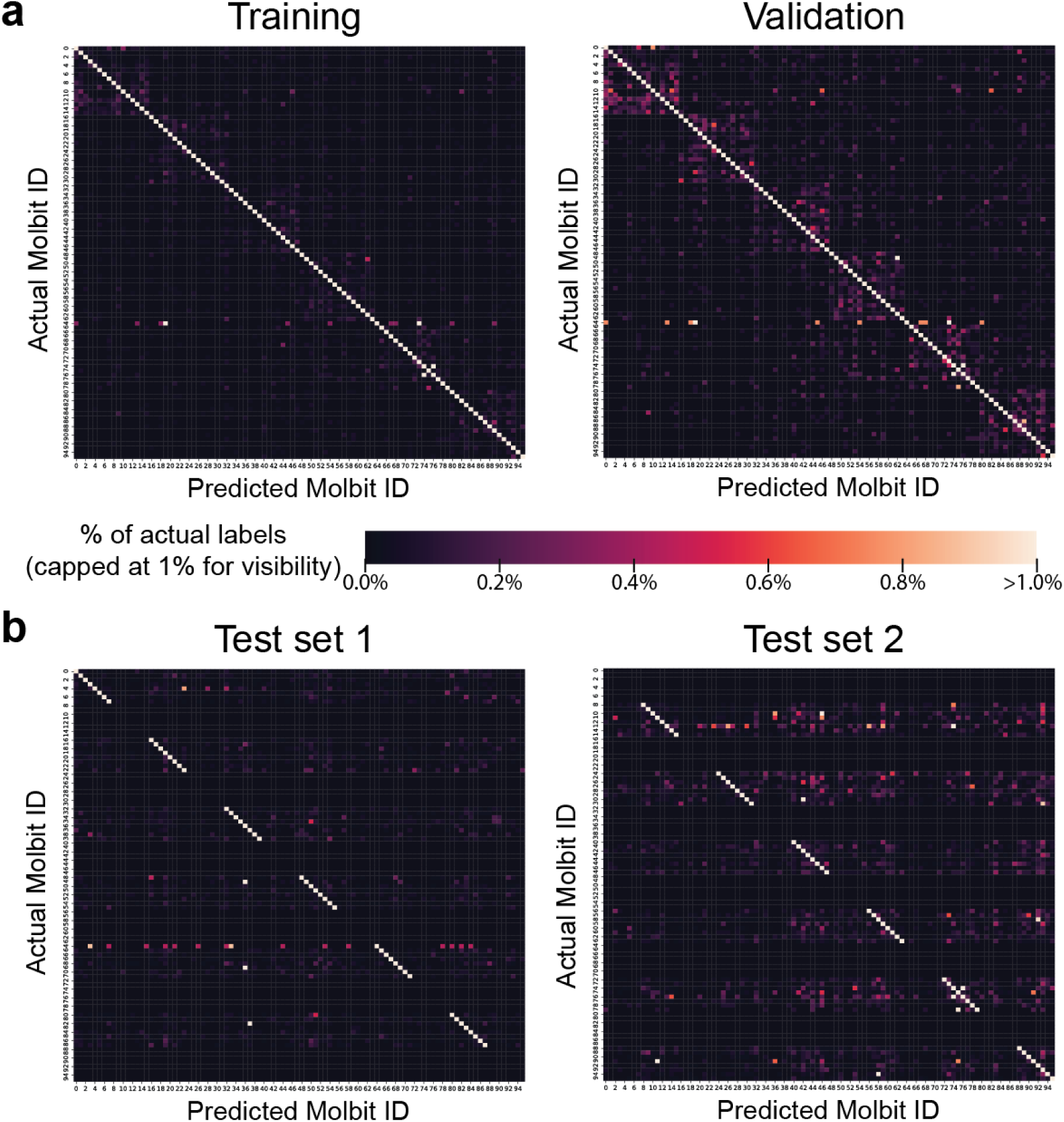
Confusion matrices for training, validating, and testing the molbit classification model. (a) Training and validation. Since counts vary drastically for each molbit, values are normalized by the total number of actual (i.e., labeled via basecalling plus alignment) molbits. Due to high overall accuracy, the visualization is capped at 1% to make error patterns more visible. In the validation plot, some batching bias is visible, demonstrated by six squares surrounding the diagonal identity line. The 16 molbits within each of these six boxes were sequenced together in the same run. (b) Testing. As in (a), values are normalized by the total number of actual molbits. Each test set consists of a mutually exclusive set of half of the molbits, in this case arranged arbitrarily in groups of 8, causing the horizontal banding in the matrices.

**Supplementary Figure 7:**
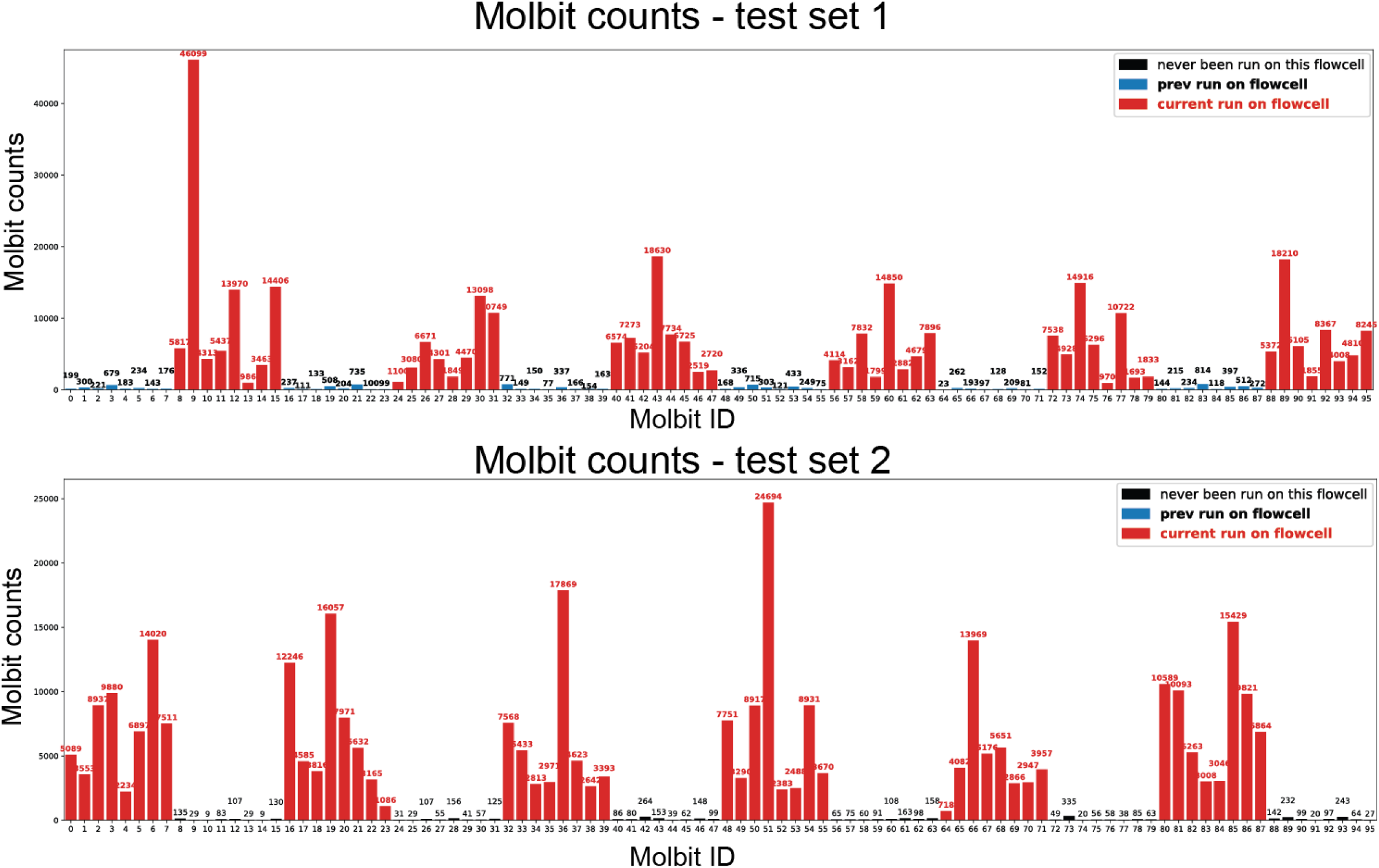
Read counts per molbit for test runs.

**Supplementary Figure 8:**
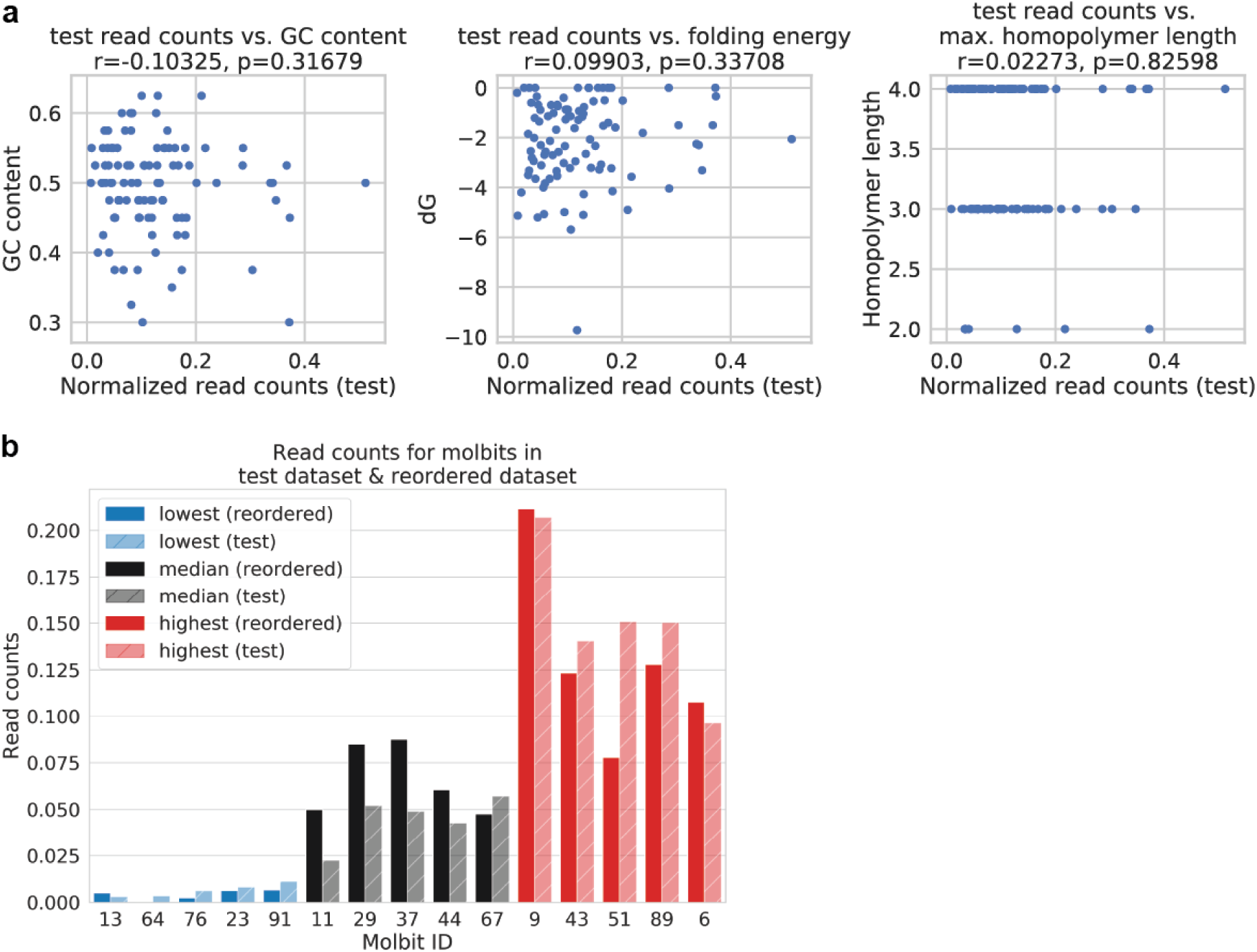
Read count variation analysis. (a) Correlation of normalized read counts against various sequence-related metrics: GC content, folding energy (*δG* minimum free energy), and maximum homopolymer length). Normalized read counts were drawn from the two datasets used to test the model, and normalized training read counts showed similar trends. (b) Normalized read counts for molbit sequences that were reordered from IDT (solid bars), compared to the same molbits in the original test datasets (hashed bars). The reordered molbits were chosen randomly out of the lowest (blue), near-median (black), and highest (red) represented molbits. Generally, molbits were detected at similar levels in both datasets.

**Supplementary Figure 9:**
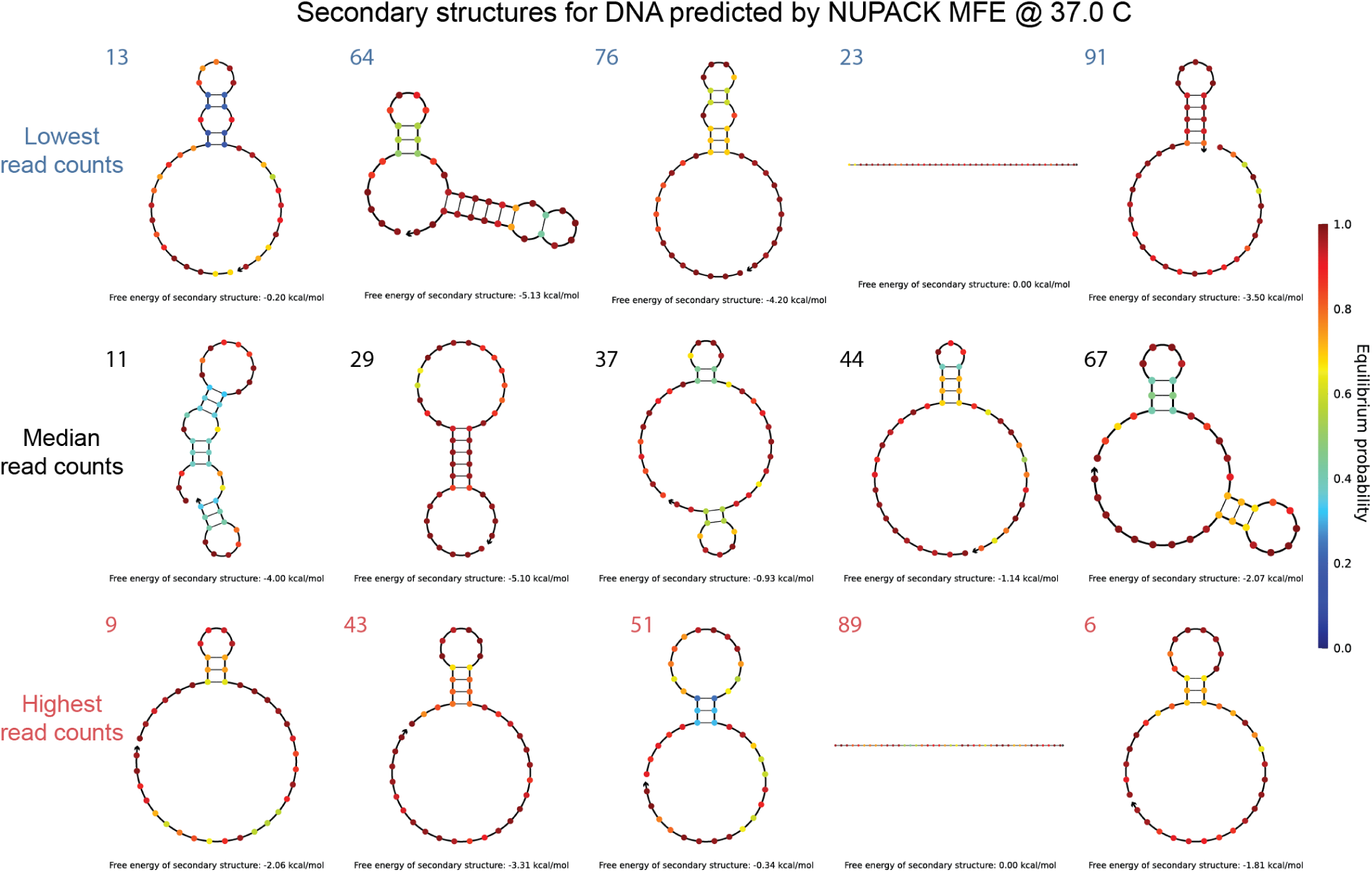
NUPACK predicted folding and minimum free energy for re-ordered sequences. Top row contains low read count sequences, middle row contains average read count sequences, and bottom row contains high read count sequences.

